# The immune landscape of primary central nervous system diffuse large B cell lymphoma

**DOI:** 10.1101/2020.08.17.254284

**Authors:** Melissa Alame, Emmanuel Cornillot, Valère Cacheux, Valérie Rigau, Valérie Costes-Martineau, Vanessa Lacheretz-Szablewski, Jacques Colinge

## Abstract

Primary central nervous system diffuse large B-cell lymphoma (PCNSL) is a rare and aggressive entity that resides in an immune-privileged site. The tumor microenvironment (TME) and the disruption of the immune surveillance influence lymphoma pathogenesis and immunotherapy resistance. Despite growing knowledge on heterogeneous therapeutic responses, no comprehensive description of the PCNSL TME is available. We investigated the immune subtypes of PCNSL and their association with molecular signaling and survival. Bulk mRNA-sequencing (n=20) and microarray (n=34) data were exploited to identify three immune subtypes of PCNSL: immune-rich, poor, and intermediate. The immune-rich subtype was associated to better survival and characterized by hyper-activation of STAT3 signaling and inflammatory signaling, *e.g.*, IFNγ and TNF-α, resembling the hot subtype described in primary testicular lymphoma and solid cancer. WNT/β-catenin, HIPPO, and NOTCH signaling were hyper-activated in the immune-poor subtype. HLA down-modulation was clearly associated with a low or intermediate immune infiltration and the absence of T-cell activation. Moreover, HLA class I down-regulation was also correlated with worse survival with implications on immune-intermediate PCNSL that frequently feature reduced HLA expression. A ligand-receptor intercellular network revealed high expression of two immune checkpoints, *i.e.*, CTLA-4/CD86 and TIM-3/LAGLS9. Immunohistopathology and digital imaging showed that TIM-3 and galectin-9 proteins were clearly upregulated in PCNSL. Altogether, our study reveals that patient stratification according to immune subtypes, HLA status, and immune checkpoint molecule quantification should be considered prior to immune checkpoint inhibitor therapy. Moreover, TIM-3 protein should be considered an axis for future therapeutic development.

## INTRODUCTION

Primary central nervous system diffuse large B-cell lymphoma (PCNSL) is a rare and aggressive extra-nodal non-Hodgkin’s lymphoma (NHL) confined to the brain, spinal cord, leptomeninges, or eyes. Newly diagnosed PCNSL accounts for 4% of brain tumors and for 4–6% of extra-nodal lymphomas in immunocompetent patients. Incidence is still increasing in elderly people [1]. The emergence of high-dose methotrexate (HD-MTX) and rituximab-based regimens in PCNSL therapy has drastically improved patient survival. However, only 50% of patients respond and 10-15% demonstrate a primary refractory disease [2], highlighting an unmet need for alternative therapeutic options.

The contribution of the tumor microenvironment (TME) to tumor aggressiveness, progression, and therapy resistance has been recognized in most tumors. By targeting the immune component, immunotherapies have revolutionized the treatment of cancers. In particular, antibodies directed against immune checkpoints (ICs) or ligands thereof, *e.g.*, PD-1/PD-L1 or CTLA-4, chimeric antigen receptor T-cells (CAR-T), and T-cell engager antibodies have demonstrated clinical benefit in B-cell malignancies [3–5]. Most notably, anti-PD-1 antibodies have been approved for therapy of relapsed/refractory classical Hodgkin’s lymphoma (HL) harboring increased PD-1+ tumor-infiltrating lymphocytes (TILs), high PD-L1 expression, and 9p21.1 *(PD-L1* gene locus) copy number alteration [6]. In this context, the exclusive brain localization of PCNSL has aroused interest for thorough TME studies.

Significant efforts have been devoted to the identification of common genetic alterations and activating oncogenic signaling in PCNSL [7–14]. These alterations mainly involve Nuclear factor-kappa B (NF-κB), B cell receptor (BCR), Toll-like receptors (TLR), Mitogen-activated protein kinase (MAPK) signaling, the DNA damage response, apoptosis, and cell cycle control. Moreover, genomic studies have suggested TME dysfunctions, such as immune evasion mechanisms, *e.g.*, *HLA* (6p21) or *B2M*(15q21.2) copy loss, *PD-L1* (9p24.1) copy number gain, and immune communication impairments, *e.g.*, *IL17REL* (22q13.33) copy gain or *IFNGR1* (6p23.3-q24) copy loss. High, intermediate, and low tumor mutational burden (TMB) were respectively identified in 19%, 71.5%, and 9.5% of patients with PCNSL in a 42-patient cohort [12], thus suggesting that anti-IC therapy could be efficient against PCNSL [15].

Multiple studies have started to unravel the main cues of the PCNSL TME, although a comprehensive description is still lacking. The presence of PD-1+ TILs and the expression of PD-L1 in either microglial cells/macrophages (tumor-associated macrophages (TAMs)) or malignant B cells were found correlated with patient outcome [12,13,16]. Some predictive biomarkers for immune checkpoint inhibitor (ICI) therapy have been identified in PCNSL, including intermediate to high TMB, 9p21.1 copy number alteration, or PD-L1 expression [12,13,16]. Nevertheless, PCNSL has so far been mainly studied as an undifferentiated entity [17] despite a manifest heterogeneity in therapeutic responses. The existence of distinct molecular subtypes has not been addressed.

We thus decided that a comprehensive study, linking a complete TME description with oncogenic signaling pathways, could help our understanding of therapy resistance and potentially uncover new therapeutic opportunities. Accordingly, we functionally characterized the immune subtypes of 54 PCNSL patient samples by combining transcriptomic data analysis with histopathology and digital imaging. We show there are three immune subtypes of PCNSL: immune-rich, poor and intermediate. We also examined the immune evasion mechanisms and the main molecular pathways associated with these different immune subtypes, highlighting new potential therapeutic opportunities, including anti-TIM-3, and the overall clinical relevance of PCNSL immune subtype classification.

## MATERIAL AND METHODS

### Patients and cohorts

Surgically resected tumors from patients with PCNSL were retrospectively retrieved from the Department of Pathology, Montpellier University Hospital (France). Snap-frozen tissues were obtained during surgery from 20 patients. The newly diagnosed patients with PCNSL provided written informed consent for tissue collection and subsequent research purposes. Patients with prior or concurrent low grade B-cell lymphoma and central nervous system (CNS) metastasis of diffuse large B-cell lymphoma (DLBCL) were excluded. This project was approved by the research ethics boards of our institution (Centre des Ressources Biologiques, CRB, Montpellier) according to the Declaration of Helsinki (AC-2010-1200 and AC-2013-2033).

To increase the size of our PCNSL cohort, we retrieved 34 additional PCNSL microarrays from Gene Expression Omnibus (GEO) (GSE34771), and 48 DLBCL transcriptomes (mRNA-sequencing) from The Cancer Genome Atlas (TCGA) to compare with PCNSL data.

### mRNA-sequencing data processing

RNA was extracted from the 20 tissue samples and quality-controlled. DNA libraries were prepared with the NEBNext Ultra II RNA-Seq kit and sequenced on a NextSeq 500 (Illumina) system using 75bp single reads. With our pipeline, Fastq files were aligned against the human genome (Ensembl GRCh38) (STAR using default parameters and 2-pass mode, read count extraction with HTSeq-count). Data are accessible from GEO (GSE155398).

### Differential gene expression analyses

Differential gene expression was analyzed with edgeR [18], imposing p-value <0.01, FDR <0.01, minimum fold-change of 2, and a minimum average of 20 (normalized) read counts for all the samples. Heatmaps were generated with ComplexHeatmap [19]. Dendrograms for the clustering of samples (columns) and genes (rows) were constructed with Ward’s method based on Euclidean distance. For color assignment, a cutoff was applied to the 2.5% highest and lowest values.

### Pathway enrichment analysis

In order to perform whole-transcriptomic and HLA status-related analysis, we implemented hypergeometric testing on differentially expressed genes to search for enriched GO terms and Reactome pathways respectively. This was followed by Benjamini-Hochberg multiple hypothesis correction with an imposed FDR <0.05, and a minimum of 5 query genes in a Reactome pathway or 3 query genes in a GO term.

Gene signatures of described signaling pathways linked to either immune response, *e.g.* gamma interferon (IFNγ) and Tumor Necrosis Factor (TNFα), or oncogenesis, *e.g.* Protein P53 (P53) and MAPK, were retrieved from the hallmark gene sets in the Molecular Signatures Database (MSigDB v7.1) [20]. Gene Set Enrichment Analysis (GSEA) was performed with the Fast GSEA Bioconductor library [20,21]. Gene signatures listed in this study were only considered as significantly enriched (adjusted p-value <0.05) when scored together with all the hallmark signatures.

### Ligand-receptor interactions and pathways

We recently defined a ligand-receptor (L-R) database [22] and a method to infer L-R interactions from bulk RNA expression [23]. The methodology originally developed for mRNA-sequencing data was adapted to microarray data by using the maximum Spearman correlation between the different probes of a given ligand and a given receptor. L-R interactions were predicted separately in the two different datasets (sequencing and microarray). In total, 673 and 393 L-R pairs were determined from microarray and RNA-sequencing data respectively. Intersection identified 165 unique high-confident L-R pairs. Brain tissue interactions were discarded, considered as *non-specific interactions*, and in all 128 L-R pairs were conserved for further analysis. Subsequently, for these a score named L-R score was computed as previously described [23].

### Histological and immunohistochemical analysis

All cases were reviewed by expert pathologists (VLS, VR, and VCM). The diagnosis of PCNSL was made on Hemataoxylin-Eosin (HE) tissue staining and was based on the WHO 2016 classification of hematopoietic and lymphoid tissue [17]. For immunohistochemical examination, 3μm-thick tissue sections from formalin-fixed paraffin-embedded (FFPE) blocks were subjected to antigen retrieval and immunostained on a Ventana Benchmark XT autostainer (Ventana Tucson, AZ, USA). The antibodies were used after appropriate antigen retrieval according to the manufacturer’s instructions (Supplementary Table 1). Association with Epstein-Barr virus (EBV) was examined by *in situ* hybridization (ISH) using Epstein-Barr encoding region (EBER). MUM1, MYC, and P53 protein expression were considered as positive if nuclear staining was observed in at least 30%, 40%, and 10% of neoplastic cells respectively. Besides PCNSL samples, we could access one *postmortem* normal brain sample for comparison. This sample was collected for autopsy purposes and processed identically by pathologists.

### Digital imaging

DAB-positive stained cells (galectin-9 and TIM-3 staining) were automatically counted using the open-source software QuPath [24]. Given both TIM-3 and galectin-9 stained tumor cells, macrophages, and endothelial cells – cell types with identifiable cell morphologies – their quantification was evaluated by counting DAB-positive pixels to improve accuracy. Intensity thresholds for pixel detection and classification were manually set for each staining type and performed identically for the all samples. For further analyses, pixel densities were estimated as the percentage of positive pixels per mm^2^ of surface area [24]. All steps were performed under the supervision of an expert pathologist (VLS). Necrosis, tissue folds, and entrapped normal structures were carefully removed.

### Interphase fluorescence *in situ* hybridization

Interphase fluorescence *in situ* hybridization (FISH) was performed on 3-μm thick tissue sections using split signal FISH DNA probes for *BCL2*/18q21 (probe Y5407; DAKO A/S), *BCL6/3q27* (probe Y5408; DAKO A/S), *MYC/*8q24 (probe Y5410; DAKO A/S), and *PDL1/*9p24.1 (PDL1, CD274 Break Apart Probe; Empire Genomics). Digital images were captured with a Metafer Slide Scanning Platform using a Leica Axioplan fluorescence microscope (Zeiss Axio Imager M1) equipped with a charge-coupled device (CCD) camera coupled to and driven by ISIS software (MetaSystem, FISH Imaging System, Germany). A total of 100 nuclei were evaluated independently by three specialists (VLS, MA, and VC). Cases were considered positive when more than 15% of the cells exhibited abnormalities in the tissue sections.

### Statistical and survival analysis

Statistical analysis was performed using the R library survminer package. Estimation of overall survival (OS) and relapse-free survival (RFS) were generated using Kaplan-Meier method. Time-to-event distributions were compared by means of a log-rank test. The dependence between clinical variables and the immune subtypes was assessed by the χ^2^ test.

## RESULTS

### Bulk transcriptomic analysis highlights contrasting immune microenvironments in PCNSL

We conducted bulk transcriptomic analysis of our PCNSL cohort (n=20) by mRNA-sequencing. Unsupervised clustering of the complete transcriptomes revealed a group of genes almost exclusively involved in immune activation (Supplementary Fig. 1, Supplementary Table 2). The expression of these genes separated tumors into three distinct groups with high, intermediate, and low immune gene expression levels (Fig. 1a). This initial observation indicated the need to examine the immune infiltrate of each tumor. In order to investigate this, we increased the size of our cohort by retrieving 34 additional PCNSL microarrays from GEO [25]. In addition, we obtained 48 DLBCL transcriptomes (mRNA-sequencing) from The Cancer Genome Atlas (TCGA). DLBCL provided a comparison of PCNSL with its nodal counterpart.

**Figure 1.**
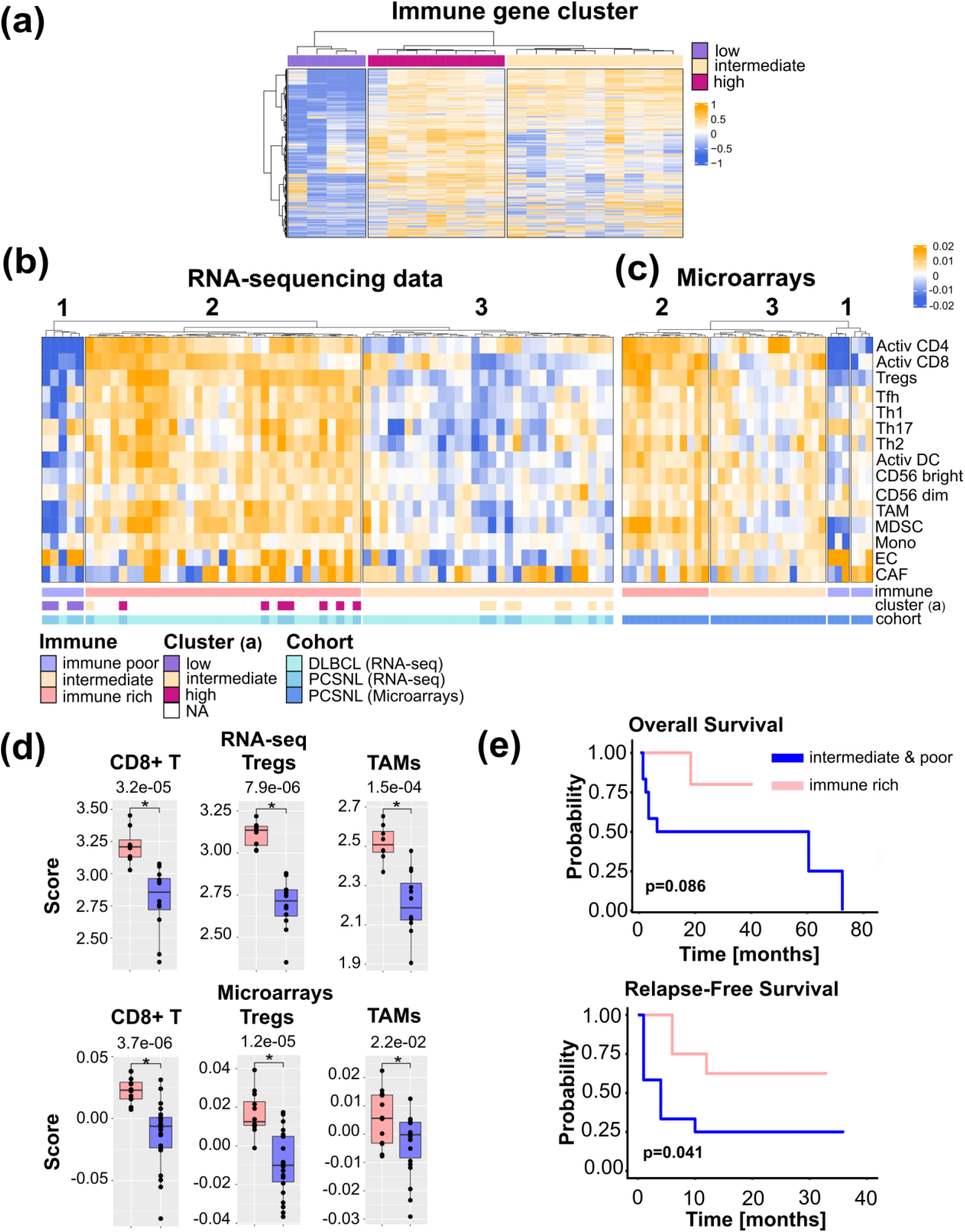
PCNSL with an immune-rich TME defines a patient subgroup with a better outcome. **a.** Three PCNSL clusters exist based on immune gene expression, denoted *high*, *intermediate*, and *low*. **b.** TME cell gene signatures across PCSNL (n=20) and DLBCL (n=48) tumor sample transcriptomes reveal three groups of tumors: the *immune-rich, immune-poor*, and *immune-intermediate* subtypes (clusters 2, 1, 3 respectively). PCNSL were classified in agreement with gene clusters in panel (a). **c.** Applying identical TME cell gene signatures to PCNSL microarray data (n=34) reveals four clusters. Merging the two small rightmost clusters yielded three groups of tumors (denoted 1-3) with immunological subtypes comparable to those of panel (b). **d.** Quantification of activated CD8+ T cells, regulatory T cells, macrophages, and CAFs in the immune-rich tumors versus the other subtype tumors (Wilcoxon one-sided tests, n=20=8+12). **e.** The immune-rich subtype features a more favorable outcome (Kaplan-Meier curve, log-rank test, n=20=8+12). Activ CD4: activated CD4+ T cells, Activ CD8: activated CD8+ T cells, Tregs: regulatory T cells, Tfh: T follicular helper cells, Th1: type 1 helper cells, Th2: type 2 helper cells, Th17: type 17 helper cells, Activ DC: activated dendritic cells, CD56 bright: CD56 bright NK cells, CD56 dim: CD56 dim NK cells, MDSC: Myeloid-derived dendritic cells, Mono: monocytes, TAM: tumor-associated macrophage, EC: endothelial cells, CAF: cancer-associated fibroblast.

We investigated 13 previously reported immune cell signatures [26] to characterize both the PCNSL and DLBCL immune infiltrates. To cover the stromal part of the TME we included signatures for cancer-associated fibroblasts (CAFs) and endothelial cells (ECs) [27]. Signature scores were computed as gene z-score averages. Based on the number of activated T cells, both mRNA-sequencing and microarray data clustered into three distinct groups (Fig. 1b-c). Considering both data types, cluster 1 (10 patients with PCNSL and one patient with DLBCL) was devoid of activated CD4+ and CD8+ T cells, but contained heterogeneous amounts of Th17 cells and macrophages. Cluster 2 (20 patients with PCNSL and 25 patients with DLBCL) was enriched in a higher number of lymphoid cells, *e.g.*, activated CD4+ and CD8+ T cells, regulatory T cells (Tregs), and myeloid cells such as TAMs, myeloid-derived suppressive cells (MDSCs), or activated dendritic cells (DCs). The last cluster, cluster 3 (24 patients with PCNSL and 22 patients with DLBCL), assumed no specific pattern with a variable immune cell presence and at a much lower level than cluster 2.

Activated CD8+ T cell, macrophage, and Treg scores were higher in cluster 2 than in clusters 1 and 3 merged (Fig. 1d). Moreover, the PCNSL classification obtained by TME cell gene signatures (Fig. 1b-c) was consistent with the groups defined in Fig. 1a. Hence, cluster 1 was termed *immune poor*, cluster 2 *immune rich*, and cluster 3 *immune intermediate.* These terms defined three immune subtypes of PCNSL. Fifty-two percent (25/48) of DLBCL tumors harbored an immune-rich phenotype versus only 37% (20/54) of PCNSL tumors. In contrast, 19% (10/54) of PCNSL tumors were devoid of CD4+ and CD8+ T cells (immune poor), whereas only one DLBCL sample (2%, 1/48) fell in this group. High proportions of both PCNSL (44%, 24/54) and DLBCL (46%, 22/48) tumors adopted an intermediate-immune phenotype.

We compared the immune-rich subtype to the other two subtypes according to overall survival (OS) and relapse-free survival (RFS). PCNSL tumors lacking an immune-rich infiltrate relapsed significantly earlier (Fig. 1e). We found a similar trend with OS; PCNSL patient tumors lacking an immune-rich infiltrate had a generally decreased OS (Fig. 1e). We did not find any association between the immune pattern and the clinical, cytogenetic, or immunohistological variables, with exception of patient outcome (Table 1). The immune phenotypes were independent of the Memorial Sloan-Kettering Cancer Center (MSKCC) score.

**Table 1.** Comparison of clinical data in the three immune groups of PCNSL

| | Patients | Immune poor<br>(n=4) | Intermediate<br>(n=8) | Immune<br>rich (n=8) | $\chi^2$ test<br>(p-value) |
| --- | --- | --- | --- | --- | --- |
| Patients |  |  |  |  |  |
| Mean age, years (range) | 64 (43-81) | 74 (62-81) | 63 (43-80) | 60 (43-70) | 0.03# |
| Male, n (%) | 10 (50%) | 2 (50%) | 4 (50%) | 4 (50%) | 1 |
| Female, n (%) | 10 (50%) | 2 (50%) | 4 (50%) | 4 (50%) |  |
| MSKCC prognostic class |  |  |  |  |  |
| Class 1, n (%) | 4 (20) | 0 | 3 (37) | 1 (13) | 0.08 |
| Class 2, n (%) | 7 (35) | 2 (50) | 0 | 5 (63) |  |
| Class 3, n (%) | 9 (40) | 2 (50) | 5 (63) | 2 (25) |  |
| Outcome |  |  |  |  |  |
| CR at last follow up, n (%) | 9 (45) | 2 (50) | 1 (12.5) | 6 (75) | 0.02* |
| AWD at last follow up, n (%) | 2 (10) | 0 (0) | 1 (13.5) | 1 (12.5) |  |
| DOD, n (%) | 9 (45) | 2 (50) | 6 (75) | 1 (13.5) |  |
| Relapse or progression, n (%) | 12 (60) | 2 (50) | 7 (88) | 3 (38) | 0.11 |
| Tumor cell phenotype |  |  |  |  |  |
| Non-GC phenotype-positive, n (%) | 15 (75) | 4 (100) | 5 (63) | 6 (75) | 0.56 |
| GC phenotype-positive, n (%) | 4 (20) | 0 (0) | 2 (25) | 2 (25) |  |
| unclassified | 1 (5) | 0 (0) | 1 (12) | 0 (0) |  |
| cMYC-positive, n (%) | 9 (45) | 2 (50) | 3 (38) | 4 (50) | 0.86 |
| BCL2-positive, n (%) | 15 (75) | 2 (50) | 7 (88) | 6 (75) | 0.38 |
| cMYC/BCL2-positive, n (%) | 8 (40) | 2 (50) | 3 (38) | 3 (38) | 0.9 |
| BCL6-positive, n (%) | 17 (85) | 4 (100) | 5 (63) | 8 (100) | 0.07 |
| CD10-positive, n (%) | 4 (20) | 0 (0) | 2 (25) | 2 (25) | 0.53 |
| EBER-positive, n (%) | 0 (0) | 0 (0) | 0 (0) | 0 (0) | 1 |
| P53-positive, n (%) | 5 (25) | 2 (50) | 0 (0) | 3 (38) | 0.09 |
| PDL1-positive, n (%) | 2 (10) | 0 (0) | 0 (0) | 2 (25) | 0.28 |
| Cytogenetic |  |  |  |  |  |
| BCL2-break positive, n (%) | 0 (0) | 0 (0) | 0 (0) | 0 (0) | 1 |
| BCL6-break positive, n (%) | 5 (25) | 0 (0) | 2 (25) | 3 (38) | 0.36 |
| cMYC-break positive, n (%) | 0 (0) | 0 (0) | 0 (0) | 0 (0) | 1 |
| PDL-break positive, n (%) | 1 (0.05) | 0 (0) | 0 (0) | 1 (13) | NA |

MSKCC: Memorial Sloan-Kettering Cancer Center; CR: complete remission; AWD: alive with disease; DOD: dead of disease; # Comparison of age of patients with immune-poor tumors to age of patients with immune-rich tumors (Wilcoxon one-sided test).

### HLA down-regulation correlates with the immune microenvironment in PCNSL

Presentation of neoantigens via HLA molecules on the surface of malignant B-cells should induce an antitumor immune response. However, B lymphoma cells can evade this response through various mechanisms, including loss or aberrant expression of HLA molecules [28]. Loss of either HLA class I or II expression has been demonstrated in PCNSL [28–31]. Nevertheless, these results remained incomplete regarding the link between the immune infiltrate and loss of HLA expression in PCNSL [28,30,32]. We thus investigated the global gene expression patterns related to HLA class I and II expression loss and potential associations with the immune subtypes identified in this study. Sixteen *HLA* genes were retrieved from the literature [33], out of which 15 were found in both PCNSL cohorts in this study, *i.e.*, in mRNA-sequencing and microarrays. For each gene, we defined low and high expression according to its median. Samples from both cohorts were pooled and hierarchical clustering of the total 54 samples identified four well-defined clusters (Fig. 2a). Seventy-eight percent (42/54) of PCNSL tumor samples featured low HLA class I or II molecule gene expression (clusters 2, 3 and 4, Fig. 2a-b), and 31% (17/54) exhibited low expressions of both (cluster 4). Twenty-two percent (12/54) featured only low HLA class I molecule gene expression (cluster 2), and lastly, 22% (12/54) had only low HLA class II molecule gene expression (cluster 3, Fig. 2a-b). Out of the 22% (13/54) of tumors featuring *HLA* expression above median expression (cluster 1), 77% (10/13) were highly infiltrated by immune cells (immune-rich subtype), whereas none of the PNCSL tumor samples with an immune-rich TME harbored *HLA* down-regulation (Fisher exact one-sided test, p-value=0.001). The HLA class I and II down-regulated cluster (cluster 4) was composed of 41% (7/17) immune-poor and 59% (10/17) immune-intermediate PCNSL tumors (Fig. 2b). Sequencing data were submitted to differential gene expression analysis between clusters 4 (HLA class I and II below median, n=8) and 1 (HLA class I and II above median, n=6) and revealed pathways related to immune activation (Fig. 2c-d, Supplementary Table 3). Moreover, patients with complete reduction of *HLA* expression relapsed earlier than patients who maintained *HLA* expression (Fig. 2e), confirming a link between HLA expression maintenance and immune-rich TME in PCNSL.

**Figure 2.**
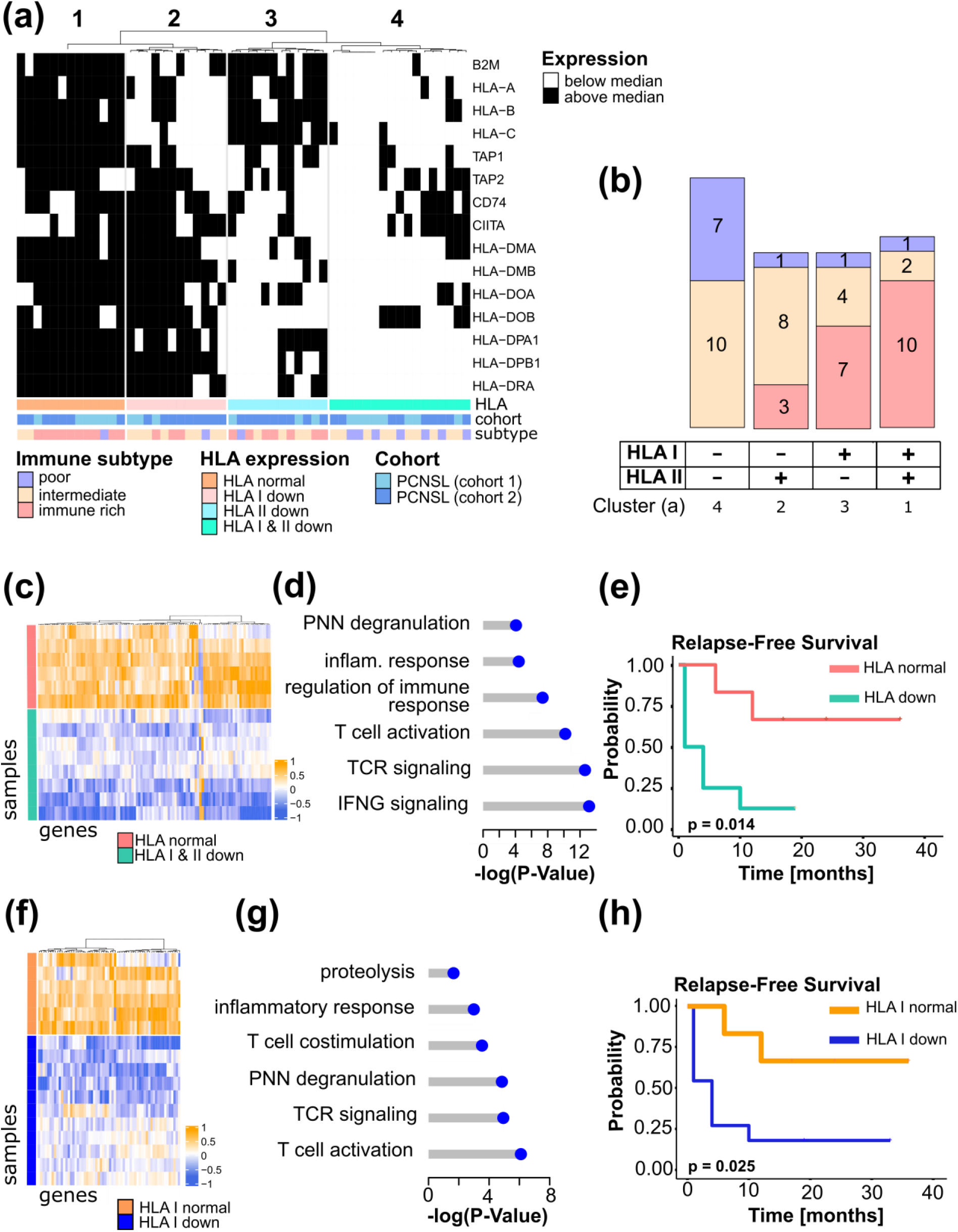
HLA expression is related to an immune-rich TME in PCNSL. **a.** HLA class I and II gene expression in PCNSL. High/low HLA gene expression defined the HLA status: HLA normal, HLA I, II, and I & II down (see heatmap legend). Expression was assessed independently in each cohort; cohort 1 is mRNA-sequencing and cohort 2 is microarray data. **b.** Samples from the distinct immune subtypes were counted in each HLA gene expression cluster of panel (a). **c.** Differentially expressed genes according to distinct HLA status (HLA normal in pink versus HLA I, II, and I & II down in green): 103 significantly deregulated genes were selected that perfectly segregated the samples according to their HLA status (FDR<0.01, log2-FC>4 (absolute value), average read counts >20). **d.** Main Gene Ontology Biological Process (GOBP) terms found significantly enriched in genes in (c) (hypergeometric test, FDR<0.05, at least 3 deregulated genes in each GO term). **e.** HLA-down status (I, II, I & II) associates with earlier relapse in PCNSL (Kaplan-Meier curves, log-rank test, n=14, normal=above median, down=below median). **f.** Differentially expressed genes between PCNSL with normal HLA class I gene expression (orange) and HLA I down (dark blue): 62 significantly deregulated genes were selected which perfectly segregated the two groups (FDR<0.01, log2-FC>4 (absolute value), average read counts >20). **g.** Main GOBP terms found significantly enriched in genes in (f) (hypergeometric test, FDR<0.05, at least 3 deregulated genes in each GO term). **h.** HLA I-down status associates with earlier relapse in PCNSL (Kaplan-Meier curves, log-rank test, n=17, normal=above median, down=below median).

We next investigated the expression of the different classes of HLA separately. HLA class I gene down-regulation was more frequent in the immune-poor than the immune-rich PCNSL tumor samples (Fisher exact one-sided test, p-value=9.10×10^-4^). Differential gene expression analysis on sequencing data between tumors with a low (cluster 2 and 4, n=11) versus higher than median (cluster 3, n=6) HLA class I gene expression revealed pathways exclusively related to immune activation, *e.g.*, costimulatory molecules, cytokines, T cell activation, and cytotoxicity (Fig. 2f-g, Supplementary Table 4). Moreover, survival analysis showed that HLA class I gene expression diminution was associated with earlier relapse in patients with PCNSL (Fig. 2h).

Dependence between HLA class II down-regulation and the immune subtypes was significant (Fisher exact one-sided test, p-value=0.025). Small homozygous deletions are known to affect the *HLA-DR* gene in PCNSL [31] and *HLA-DRA* expression was reported as a prognostic factor in DLBCL [34]. We hence compared *HLA-DRA* expression between the three immune subtypes and found it more highly expressed in the immune-rich group than in the immune-poor PCNSL (Fisher exact one-sided test, p-value=6.10^-4^) or the immune-intermediate (Fisher exact one-sided test, p-value=0.01) tumors. In particular, *HLA-DRA* expression was correlated with activated CD8+ T cells (spearman correlation r=0.71, p-value=4.10^-4^) and Th1 cells (spearman correlation r=0.53, p-value=0.0027). Differentially expressed genes between *HLA-DRA* normal versus loss of gene expression samples identified genes related to the innate immune response, cytokines, antigen presentation, and extracellular matrix (ECM) organization (Supplementary Fig. 2a-b, Supplementary Table 5). Contrary to patients with DLBCL, patients with PCNSL showed no association between *HLA-DRA* gene expression and outcome (Supplementary Fig. 2c-d).

### Immune subtypes of PCNSL adopt distinct oncogenic signaling

Next, we explored the potential relations between the PCNSL immune subtypes identified in our present study and the deregulated oncogenic pathways discussed in the literature on PCNSL [7–9,11–14,35,36]. The NF-κB, Signal transducer and activator of transcription 3 (STAT3), IFNγ, Phosphoinositide 3-kinase/Protein kinase B (PI3K/AKT), Kirsten ras sarcoma viral oncogene (KRAS), Vascular endothelial growth factor (VEGF), P53, MAPK, Salvador-Warts-Hippo (HIPPO), Interleukin-10 (IL-10), TNF-α, WNT/β-catenin, Transforming growth factor beta (TFG-β), and NOTCH pathways were investigated. We retrieved their gene signatures from the Molecular Signatures DataBase (MSigDB) [20]. Average z-scores of signature genes were computed as previously described [37].

We observed distinct activations of the oncogenic pathways depending on the immune subtypes of PCNSL (Fig. 3a). The immune-rich tumors logically featured strong activation of signaling pathways known to be involved in the immune response, *e.g.*, STAT3, IFNγ, IL-10, TNF-α, and NF-κB, as well as in stromal signaling, *i.e.*, TGF-β. Activation of KRAS and P53 signaling appeared also favored in immune-rich PCNSL samples, although in a less pronounced manner and not exclusively. In contrast, WNT/β-catenin, NOTCH, and HIPPO pathways were more activated in the immune-poor PCNSL samples. GSEA was performed to confirm the differences observed between the immune-rich and the other PCNSL subtypes (Fig. 3b). Signature scores for NOTCH, WNT/β-catenin, and HIPPO for each immune subtype are featured in Fig. 3c.

**Figure 3.**
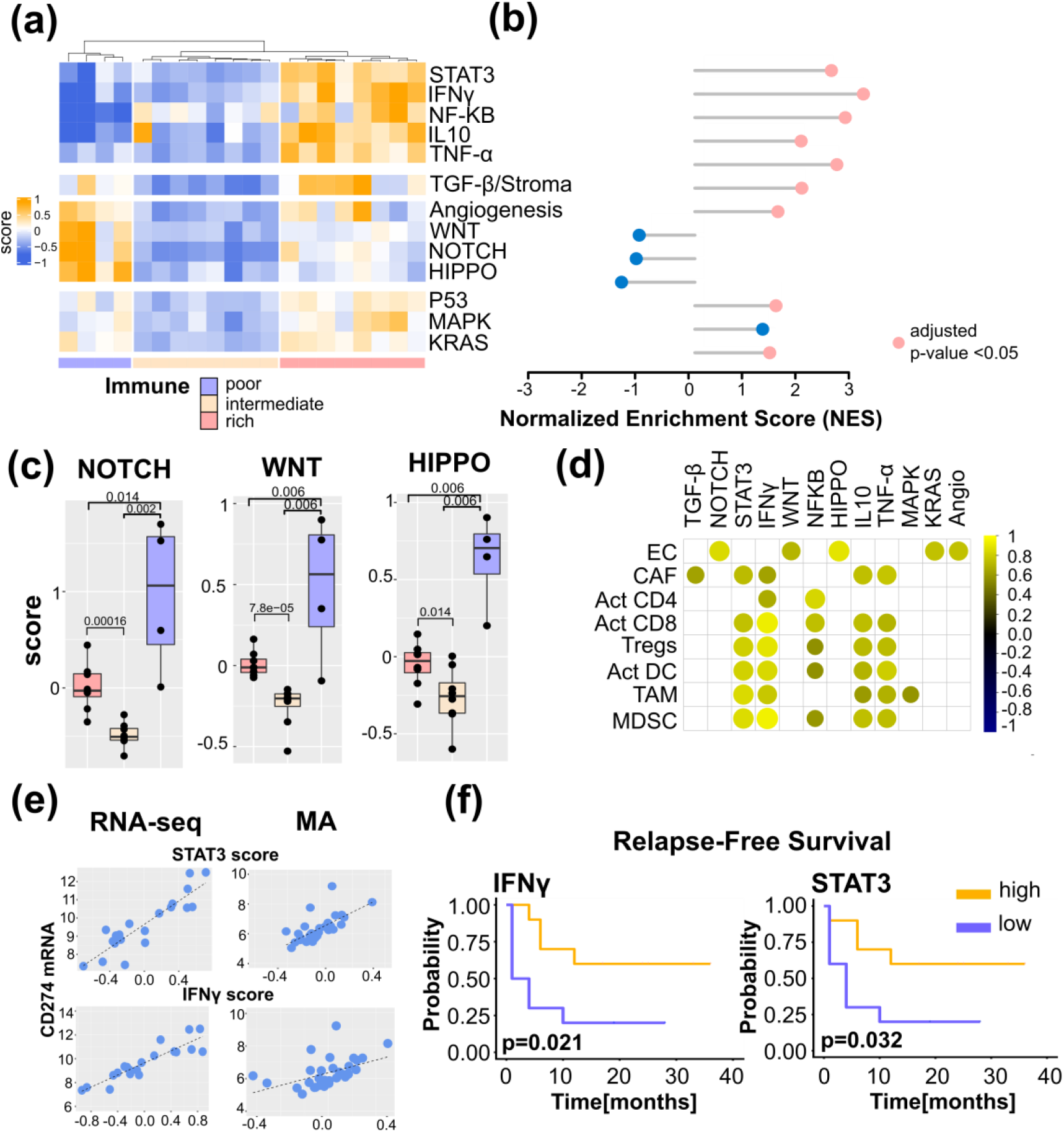
The immune landscapes of PCNSL unravel distinct oncogenic signaling. **a.** Signature scores (average z-scores of signature genes) of different oncogenic signaling pathways according to the immune subtypes defined in Fig. 1bc. **b.** Gene Set Enrichment Analysis performed between the immune-rich and other subtypes (intermediate & poor). We report the normalized enrichment score (NES) and indicate statistical significance (FDR<0.05) by a pink dot (otherwise blue). **c.** Quantification of NOTCH, WNT, and HIPPO signature scores in the different immune subtypes (Wilcoxon one-sided tests, p-value <0.05, n=4+8+8=20, one immune-poor subtype outlier was removed (significant according to Grubbs and Dixon tests). **d.** Correlation between the main cell-type scores (EC, CAF, Act CD8, Tregs, Act DC, TAM, MDSC) and oncogenic signaling z-scores. Highly significant correlations are indicated only (spearman rank correlation coefficient, r>0.6, p-value <0.01). **e.** Correlation between STAT3 and IFNγ signature scores and CD274 gene expression in the PCNSL cohort (spearman rank correlation coefficient, r>0.6, p-value <0.0001). Note that correlation is observed with both mRNA-sequencing (RNA-seq) and microarray (MA) data. **f.** High INFγ and STAT3 signaling z-scores were associated with lower relapse-free survival (RFS) in PCNSL (Kaplan-Meier curves, log-rank test, n=20, high=above median, low=below median).

We then assessed correlations between the main cell types we used to profile PCNSL TME (Fig. 1b-c), *i.e.*, activated CD8+ T cells, activated CD4+ T cells, Tregs, macrophages, DC, EC, and CAF, and the different signaling pathways (Fig. 3d). The TGF-β z-score was correlated with CAF abundance as expected, while the four immune cell types (activated CD8+ T cells, Tregs, TAMs, and DCs) were highly correlated with IFNγ, TNF-α, IL-10, and STAT3 z-scores. Angiogenesis, as well as WNT/β-catenin, NOTCH, and HIPPO pathways correlated with the EC signature score.

STAT3 and IFNγ signaling pathways are generally considered coupled to PD-L1 up-regulation in solid tumors and DLBCL [38]. Although a consensual good prognostic value for IFNγ activity has been acknowledged, hyper-activation of STAT3 signaling has been associated with varying survival outcomes [38–40]. Given a link between PD-L1-expressing TAMs and STAT3 signaling has been reported in PCNSL [41], we exploited a PCNSL retrospective cohort of 57 patients [16] that we previously described in the assessment of an association between high PD-L1-expressing TAMs and survival. We defined two histological scores for this purpose: TAM density, evaluated by CD68-positive staining, and the percentage of PD-L1-positive TAMs. High PD-L1 protein expression in TAMs was defined as the simultaneous detection of high TAM density (grades 2-3) and high PD-L1 expression in these TAMs (>50% of CD68-positive staining). High PD-L1-expressing TAMs were found to be strongly associated with better survival and a lower risk of relapse in patients with PCNSL (Supplementary Fig. 3a). At the gene expression level, the PD-L1 transcript *(CD274)* was highly correlated with STAT3 and IFNγ signature scores in RNA-sequencing and microarray data (Fig. 3e). Segregating patients according to STAT3 and IFNγ median expressions resulted in an association of high STAT3 and IFNγ expressions with lower relapse risk (Fig. 3f). We also observed a trend with OS but not a significant association (Supplementary Fig. 3b-c).

### Mapping PCNSL intercellular interactions

Intercellular interactions, particularly ICs, within the TME are known to contribute to tumor progression and therapy resistance. In this context, specific ligand-receptor (L-R) interactions, *e.g.*, PD-1/PD-L1, have been extensively studied in solid tumors and PCNSL [12,16,41,42]. We recently proposed an algorithm to infer L-R interactions from bulk transcriptomics [23]. This algorithm was applied in this study separately to mRNA-sequencing and microarray data (Fig. 4a, Supplementary Tables 6-7). This resulted in the identification of 165 confident L-R pairs, from which brain cell-related interactions were discarded as potential background noise, yielding a total of 128 PCNSL-specific confident L-R pairs (Supplementary Table 8). Since our algorithm associates each L-R pair to signaling pathways in order to test receptor downstream activity, we summarized recurrent pathways in Fig. 4b. These pathways largely reflect the oncogenic signaling pathways already investigated above in Fig. 3. We then searched for immune infiltration-related L-R pairs by computing a score (the L-R score), indicative of the L-R pair co-expression level in a given sample [23], and correlated that score with immune infiltration (sum of all the immune cell signatures) as reported in Fig. 1. We found 46 correlated L-R pairs (Spearman correlation, r>0.5, adjusted p-value<0.05, Fig. 4c). Among these, we denoted several ICs, *e.g.* CD86/CTLA4, LGALS9/HAVCR2, LILRB2 and its ligands, as well as inflammatory pairs, *e.g.*, B2M/CD247, and CCR5 and its ligands. Other pairs, *e.g.*, ANGPT1/TEK, DLL1/NOTCH1, THBS1/ITGB1, were linked to angiogenesis according to the pathways used by our algorithm. Some L-R pairs, *e.g.*, SELPLG/ITGB2, PDGFB/LRP1, C3/C3AR1, and LGALS9/HAVCR2, were expressed in PCNSL devoid of lymphoid cells (Fig.4c).

**Figure 4.**
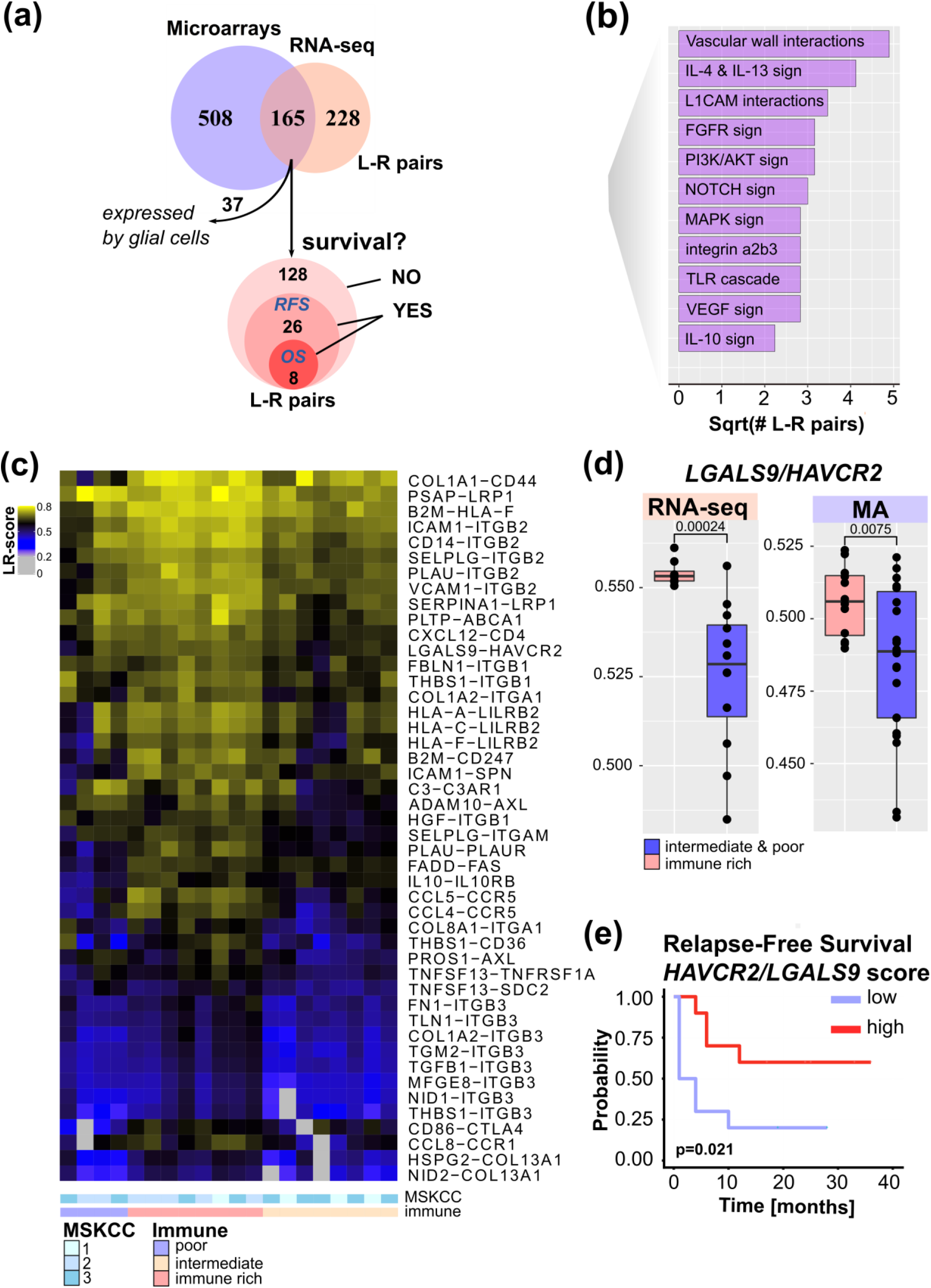
Ligand-receptor interactions within the PCNSL microenvironment. a. Ligand-receptor (L-R) pair selection strategy: our algorithm selected 673 and 393 confident L-R pairs from the mRNA-sequencing (n=20) and microarray (n=34) datasets respectively. A total of 128 confident and PCNSL-specific L-R pairs were selected from both datasets. Out of these 128 L-R pairs, survival analysis found 26 L-R pairs significantly associated to RFS and 8 to overall survival (OS). b. Functional categories associated to the L-R pairs selected. c. Immune infiltrate-associated L-R pairs in our PCNSL cohort (n=20). d. *LGALS9/HAVCR2* L-R scores are higher in the immune-rich subtype of PCNSL (Wilcoxon one-sided tests, p-value <0.01, n=20=8+12 and n= 2+22 for mRNA-sequencing and microarrays data respectively) e. *LGALS9/HAVCR2* above median L-R scores are associated with better RFS (Kaplan-Meier curve, log-rank test, p-value <0.05).

Among the 128 PCNSL-specific confident L-R pairs, we found 26 pairs significantly associated with RFS. These included eight pairs significantly associated with both OS and RFS (Fig. 4a and Supplementary Table 8). *LGALS9/HAVCR2* L-R scores were higher in the immune-rich PCNSL samples (Fig. 4d) and were significantly associated with RFS (Fig. 4e). T-cell immunoglobulin mucin receptor 3 (TIM-3, *HAVCR2* gene) is an IC receptor that mainly plays a role in T-cell exhaustion. It suppresses T cell responses upon its binding to galectin-9 *(LGALS9* gene). However, TIM-3 has demonstrated several behaviors depending on context [43], and hence its role in the brain TME of PCNSL demands investigation. Protein expression of TIM-3 and its ligand, galectin-9, was quantified by digital imaging in 32 PCNSL patient tumor samples from a retrospective cohort we published recently [16] and in one *postmortem* normal brain sample (Fig. 5a-b). In the normal brain tissue, we observed Galectin-9 protein expression in glial cell ramification, ECs, and rare macrophages, while TIM-3 was rarely expressed (only by a very few ECs and macrophages) (Fig. 5a). In the tumors, we found up-regulation of both TIM-3 and galectin-9 expression (Fig. 5b-c). TIM-3 and galectin-9 protein expression were also correlated in tumor samples (Fig. 5d). TIM-3 was mainly expressed by tumor cells, TAMs, and small lymphocytes, whereas galectin-9 was mainly expressed by TAMs, ECs, glial cells, and gemistocytes. Notably, galectin-9 was strongly expressed in the tumoral area and glia, characterized areas of brain inflammation. TIM-3 expression increased with increasing MSKCC score, whereas galectin-9 seemed to be higher in both MSKCC scores 2 and 3 compared to score 1, while not demonstrating any significant difference between scores 2 and 3 (Fig. 5e). We also observed that higher TIM-3 and galectin-9 expression tended to be associated with relapse (Fig. 5f), thus in line with transcriptomic data (Fig. 4e). Given the clinical relevance of TIM-3/galectin-9, we looked at its role within the TME of PNCSL. TIM-3 (*HAVCR2*) and galectin-9 (*LGALS9*) gene expression were highly correlated with most IC ligands and/or receptors, *e.g.*, HVEM, HVEML, LAG3, PD-L1, IDO1, and CD86 (Fig. 5g). We also found that *LGALS9* and *HAVCR2* gene expressions were highly correlated with gene signatures of cell types that were enriched in the immune-rich subtype of PCNSL, *i.e.*, TAMs, Th1, MDSC, and TFh (Fig. 5h). Finally, *LGALS9* and *HAVCR2* expression was highly correlated with STAT3 and IFNγ z-scores, and correlated well with TNF-α, MAPK, and IL-10 signaling. *HAVCR2* gene expression was also correlated with TGF-β and P53 (Fig. 5i).

**Figure 5.**
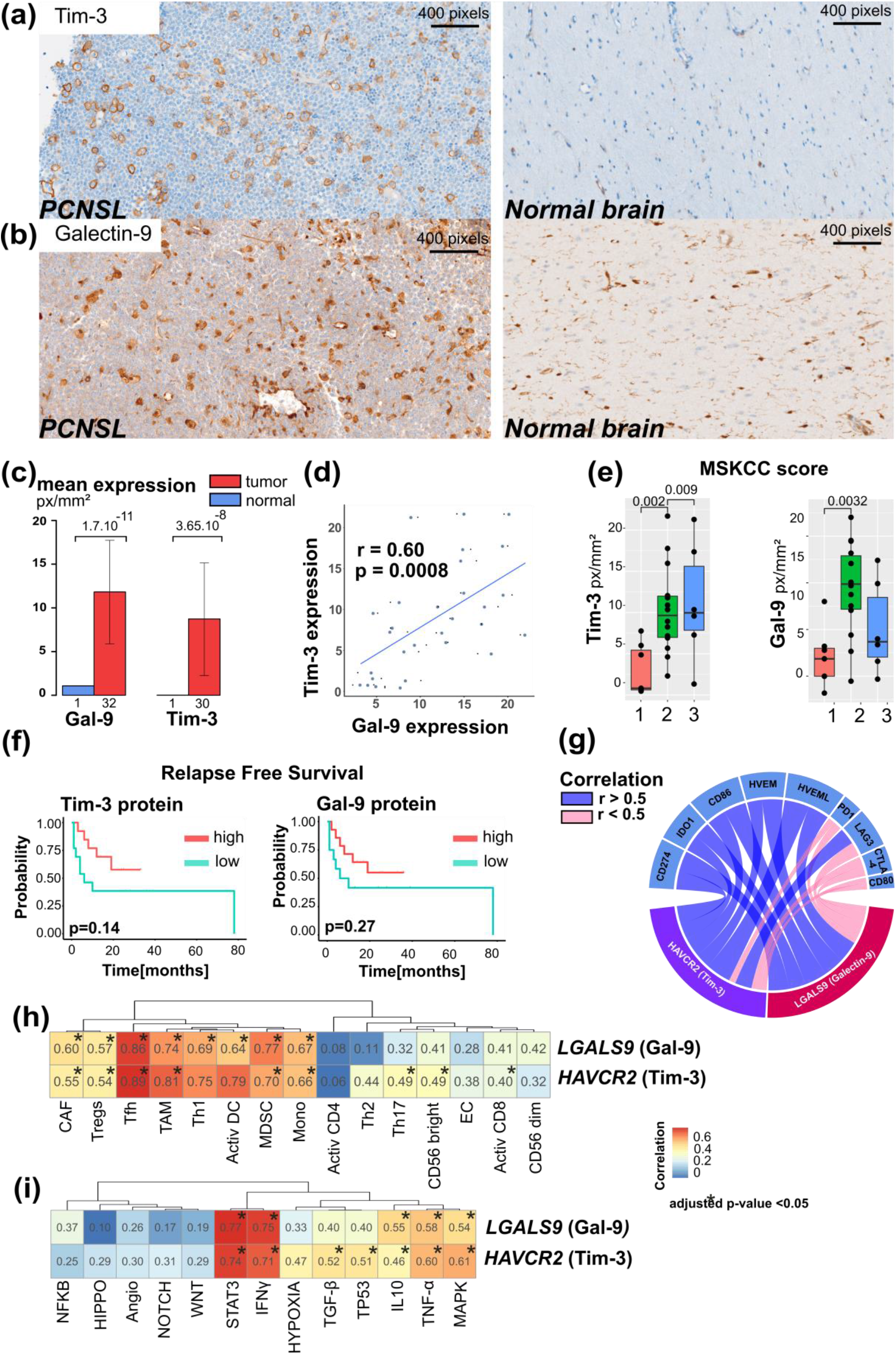
Galectin-9/TIM-3 crosstalk is up-regulated and linked to immune activities in PCNSL. a. TIM-3 protein expression is up-regulated in PCNSL compared to normal brain tissue. b. Galectin-9 (Gal-9) protein expression is up-regulated in PCNSL tissue compared to normal brain tissue. c. TIM-3 and galectin-9 expression are up-regulated in PCNSL tumors in comparison to normal brain tissue (student test). d. TIM-3 and galectin-9 expression are correlated in our PCNSL cohort (spearman rank correlation, n=20). e. TIM-3 expression is associated with MSKCC scores (1, 2, and 3), while galectin-9 only features low expression in MSKCC score 1 samples (Wilcoxon one-sided tests). f. Lower TIM-3 and galectin-9 protein expression tends to be associated with earlier relapse in PCNSL (Kaplan-Meier curves, log-rank test, n=27, high=above median, low=below median). g. Correlation between either *HAVCR2* (TIM-3) or *LGALS9* (galectin-9) gene expression and other immune checkpoints (spearman rank correlation, adjusted p-value <0.01). h. Correlation between either *HAVCR2* (TIM-3) or *LGALS9* (galectin-9) expression and the cell-type z-scores (spearman rank correlation, adjusted p-value <0.05). i. Correlation between either *HAVCR2* (TIM-3) or *LGALS9* (galectin-9) gene expression and oncogenic signaling pathway z-scores (spearman rank correlation, adjusted p-value <0.05).

## DISCUSSION

Bulk transcriptomic analysis of PCNSL patient samples allowed us to recognize three well-defined subtypes related to the immune infiltrate abundance and composition. We defined these as immune-rich, intermediate, and poor subtypes. These subtypes were independent of MSKCC score, suggesting that the immune reaction was independent of the disease stage. Moreover, comparing PCNSL and DLBCL, we found immune-poor tumors enriched in PCNSL, whereas DLBCL tumors were relatively more abundant in the immune-rich subtype. This indicates that the particular brain microenvironment could influence the immune response in PCNSL physiopathology.

We found specific associations when correlating the three PCNSL immune subtypes with oncogenic pathways commonly deregulated in this entity [7–9,11–14,35,36]. Indeed, inflammatory pathways, *e.g.*, IFNγ or NF-κB, as well as anti-inflammatory pathways, *e.g.*, STAT3 or TNF*α*, were found active in immune-rich PCNSL. In addition, IL-10, TGF-β signaling, and angiogenesis mediated by VEGF are all STAT3 activators [44] and were also found up-regulated in the immune-rich PCNSL. Immune-poor PCNSL harbored activated WNT/β-catenin, HIPPO, and NOTCH signaling. All the deregulated pathways in the immune-poor subtype were correlated with ECs and are known to play a role in angiogenesis [45,46]. No clear association was found with the immune-intermediate subtype, which is not surprising given its heterogeneous immune cell composition.

Pathways related to ICs are of particular interest to tumor biology and the clinics. The PD-1/PD-L1 interaction has been described in PCNSL [16,41] and PD-L1 was found to be produced by TAMs [16] in this tumor. We show here that high PD-L1-expressing TAMs were associated with better survival and lower relapse. Interestingly, we also showed that two important PD-1/PD-L1-regulated mechanisms (STAT3 and IFNγ signaling) were active in the immune-rich subtype and were associated with improved survival. This highlights the prognostic value of PD-L1-expressing TAMs in PCNSL.

Another IC interaction, *LGALS9* (galectin-9)/*HAVCR2* (TIM-3), was identified by a new algorithm that we recently described [23]. High *LGALS9/HAVCR2* co-expression scores (the so-called L-R scores) were found in every immune subtype of PCNSL, although they were higher in the immune-rich tumors. Concomitant up-outcome and prognosticregulation of TIM-3 and galectin-9 proteins occurred in PCNSL tumors compared to normal brain tissue, an observation already reported for other tumors in the brain [47–49]. In contrast to PD-1, we show that TIM-3 was mostly produced by both tumor cells and TAMs in PCNSL, an observation already reported for DLBCL [50], whereas it is mainly produced by CD8+ T cells and microglial cells in brain tumors, such as glioma [47]. This result suggests different intercellular communication and immune surveillance escape mechanisms in PCNSL compared to solid brain tumors. *HAVCR2* gene expression was associated to the presence of other IC molecules, inflammatory pathways, *e.g.*, STAT3 or IFNγ, and the abundance of immune cells found in the immune-rich subtype of PCNSL. Previous reports have suggested a regulatory role of TIM-3 on the expression of other IC molecules, such as PD-L1 in glioma [47], thus implying that anti-TIM-3 agents could be used to strengthen anti-PD-1 agents. Besides, in our present study high TIM-3 protein expression was associated with high MSKCC scores, and could thus serve as marker of bad prognosis in PCNSL.

The immune-poor subtype of PCNSL featured low HLA class I and II molecule gene expression, whereas immune-rich PCNSL associated with higher *HLA* expression. The immune-intermediate subtype predominantly harbored low expression of HLA class I or both class I and II genes. Low HLA class I gene expression was associated with low immune infiltration, an absence of T-cell activation, and earlier relapse, in all corroborating that low HLA class I gene expression results in poor prognosis in solid cancers [51,52]. This observation is also in agreement with the bad prognosis linked to low immune infiltration we observed in PCNSL. In our study, *HLA DRA* gene down-regulation was associated with antigen presentation, innate and adaptive immune responses, activated CD8+ T cells and Th1 cells. Although *HLA-DRA* gene expression was recognized as an independent adverse prognosis factor for RFS in DLBCL [53], we did not find any association with patient outcome in PCNSL, a result that may be linked to the poorer overall outcome of patients with PCNSL compared to DLBCL [54].

Altogether, our results highlight the similarities between the immune-rich, intermediate, and poor subtypes of PCNSL and the hot, intermediate, and cold subtypes observed in primary testicular lymphoma (PTL) [55] and in solid tumors [56]. Despite the obvious differences in terms of spatial architecture between PCNSL and solid tumors, analogies could be drawn between PCNSL immune-intermediate tumors displaying reduced HLA gene expression and solid and altered tumors [57,58]. Herein, knowledge on these PCNSL immune subtypes could help in stratifying patients prior to treatment selection. Most of our patients (18/20) were treated with immunochemotherapy containing rituximab. Despite the modest size of our cohort due to the rarity of PCNSL, we did however show longer RFS for patients harboring an immune-rich tumor and preserved HLA class I and II gene expression. This result emphasizes the informative value of PCNSL immune patterns. One limitation of our study remains the lack of a validation cohort. Nonetheless, a T cell-inflamed signature has been found associated to favorable patient outcome in a PTL cohort treated with a rituximab-containing immunochemotherapy [55].

Several PCNSL clinical trials with ICIs are ongoing. Despite a high response rate in HL, melanoma, and lung cancers [59,60], response to ICIs remains generally heterogeneous [61]. The status of HLA machinery [62], as well as the presence of active signaling pathways (such as STAT3), are considered as immune evasion mechanisms that influence responses to ICIs [63]. Therefore, patients with PCNSL harboring the immune-rich subtype could be further stratified by assessing potential hyper-activation of STAT3 signaling. In such a case, a combined regimen including an ICI and a STAT3 inhibitor could be envisioned [64]. Patients with immune-poor PCNSL are obviously less likely to benefit from ICI. Tumors featuring an intermediate-immune subtype and down-regulation of either HLA class I or II might not respond optimally to ICI monotherapy. Nevertheless, novel immunotherapies that have emerged in lymphoma, such as targeting CAR-T cells, cancer vaccines, or bispecific antibodies [3–5] could help in restoring an immune response in such intermediary immune subtypes.

NF-κB signaling was also found hyper-activated in immune-rich PCNSL. Previous studies have shown that *MYD88* gene mutations are highly prevalent and support lymphoma growth through NF-κB signaling in PCNSL [10–13]. It would hence be reasonable to co-assess the genomic and immune phenotypic statuses of PCNSL to propose combined ibrutinib/ICI therapies to immune rich/*MYD88*-mutated PCNSL patients (ongoing trial NCT03770416).

Finally, reported epigenetic profiles have revealed different methylation profiles among patients with PCNSL [65]. In particular, the *PTPN6* gene promoter region was found highly methylated in 48.5% of PCNSL tumors, leading to STAT3 hyper-activation [66]. Given that epigenetic regulators can cross the blood-brain barrier (BBB), such as DNA methyltransferase (DNMT) inhibitors, one might hypothesize that DNMT inhibitors could restore a normal STAT3 expression in immune-infiltrated PCNSL tissues. For immune-poor PCNSL, another option would be to re-establish the HLA class II expression in lymphoma cells by the use of histone deacetylase (HDAC) inhibitors [67] that can cross the BBB. These speculations further highlight the profound consequences of PCNSL immune subtype stratification, for instance in the addressing of immune-intermediate tumors.

In conclusion, we characterized the immune landscape of human PCNSL by combining bulk transcriptomic analysis, histopathology, and digital imaging. The immune-rich subtype is associated with HLA expression preservation, activation of specific signaling, *e.g.*, IFNγ, STAT3, or NF-κB, and expression of inhibitory ICs, such as PD-1/PD-L1 and TIM-3/Galectin-9. The immune-poor subtype of PCNSL is characterized by active WNT/β-catenin, NOTCH, and HIPPO signaling, limited presence of active T-cells, and down-regulation of HLA expression. Several immune evasion mechanisms and new potential therapeutic opportunities, including anti-TIM-3, highlight the clinical relevance of PCNSL immune subtype classification.

## Acknowledgements

We thank the Centre des Ressourses Biologiques of Montpellier University Hospital for access to tissue samples. We also thank Dr Simon Cabello-Aguilar for help preparing this manuscript. MA was supported by a GIRCI SOOM API-K 2016-811-DRC-AC grant, and JC was supported by Fondation ARC PJA 20141201975 and Labex EpiGenMed ANR 10-LABX-0012 grants.

