## Supplementary Material for "The immune landscape of primary central nervous system diffuse large B cell lymphoma"

[1] Institut de Recherche en Cancérologie de Montpellier (IRCM), INSERM, Parc Euromédecine, 208 rue des Apothicaires, 34298 Montpellier, France
[2] Biological Hematology Department, CHU Montpellier, Hôpital Saint Eloi, 34275 Montpellier, France
[3] Université de Montpellier, Faculté de Pharmacie, 15 avenue Charles Flahault, 34093 Montpellier, France
[4] Institut régional du Cancer Montpellier (ICM), Parc Euromédecine, 208 rue des Apothicaires, 34298 Montpellier, France
[5] Université de Montpellier, Faculté de Médecine, 2 rue école de Médecine, 34060 Montpellier, France
[6] Département de pathologie et d'oncologie biologique, CHU Montpellier, Hôpital Gui De Chauliac, 34000 Montpellier, France
[7] Equal contribution

### Supplementary Tables

**Supplementary Table 1. References and dilutions of primary antibodies used for IHC.**

| Target | Manufacturer | Cat. No. | Dilution IHC |
|---|---|---|---|
| CD20 | Dako Denmark A/S | L26 | 1/300 |
| CD3 | Ventana, PREP Kit Ventana | 2GV6 | undiluted |
| CD5 | Dako, Denmark A/S | 4C7 | 1/100 |
| CD4 | Ventana PREP Kit Ventan | SP35 | undiluted |
| CD8 | Ventana, PREP Kit Ventana | SP57 | undiluted |
| CD10 | Menarini, California USA | 56C6 | 1/10 |
| BCL6 | Ventana PREP Kit Ventana | G1191E/A8 | undiluted |
| MUM1 | Dako | MUM1p | 1/50 |
| P53 | Ventana, PREP Kit Ventana | DO7 | undiluted |
| MYC | Epitomics, Burlingame, CA, USA | EP 121 | 1/100 |
| CD68 | DAKO | KP1 | 1/400 |
| CD163 | Ventana, PREP Kit Ventana | MRQ-26 | undiluted |
| KI67 | Ventana, PREP Kit Ventana | 30-9 | undiluted |
| PD1 | Abcam, Paris FRANCE | NAT105 | 1/100 |
| PDL1 | Cell Signalling Technology | E1L3M | 1/200 |
| TIM3 | Cell Signalling Technology | D5D5R | 1/100 |
| Galectin-9 | OriGene Technologies | OTI8B11 | 1/150 |

**Supplementary Table 2. GO terms enriched in the 4 gene clusters**

**Supplementary Table 3. Pathways (GO terms and Reactome pathways) associated with HLA status in PCNSL**

**Supplementary Table 4. Pathways (GO terms and Reactome pathways) associated with HLA class I gene expression in PCNSL**

**Supplementary Table 5. GO terms associated with high *HLA-DRA* gene expression in PCNSL**

**Supplementary Table 6. L-R pairs selected by RNA-sequencing analysis (n=20)**

**Supplementary Table 7. L-R pairs selected by microarrays analysis (n=34)**

**Supplementary Table 8. L-R pairs selection**

### Supplementary Figures

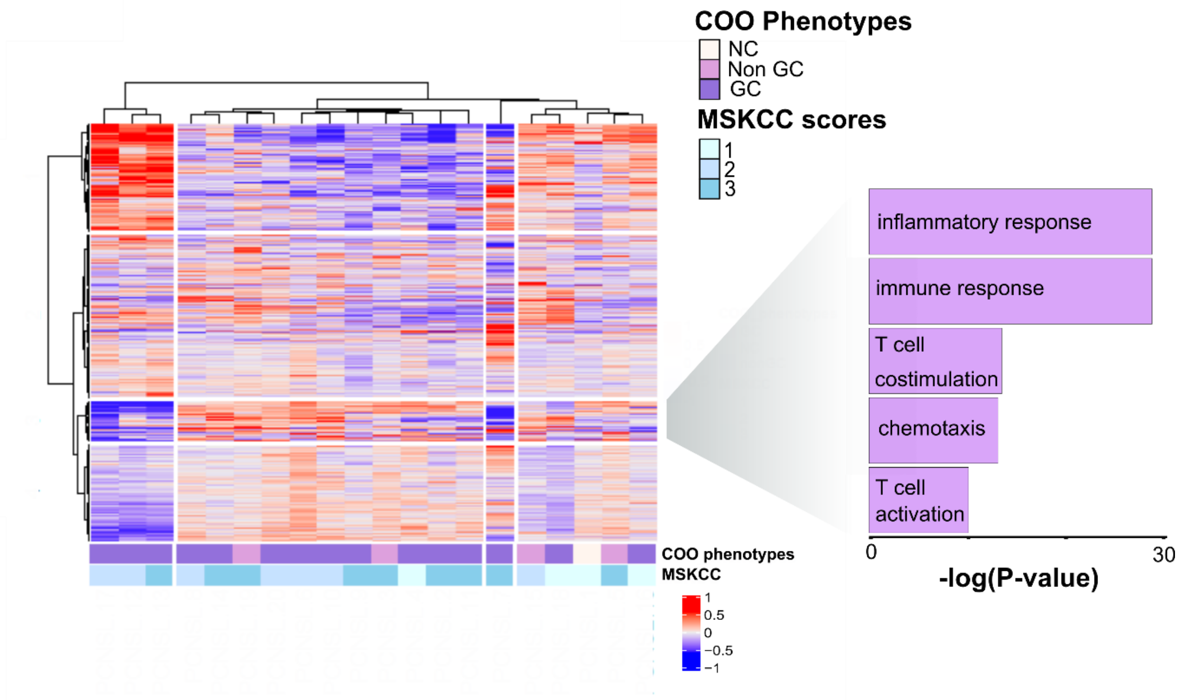


**Supplementary Fig. 1 An immune gene expression cluster revealed heterogeneity among PCNSL samples**

Four clusters of genes were revealed by unsupervised clustering of our 20 PCNSL complete transcriptomes (Ward’s method based on Euclidean distance). The main GOBP terms found enriched in cluster 3 were almost exclusively related to immune activation (hypergeometric test, FDR <0.01, at least 10 deregulated genes in each GO term to limit the size of the figure). The full list of GOBP terms is provided in Supplementary Table S2.

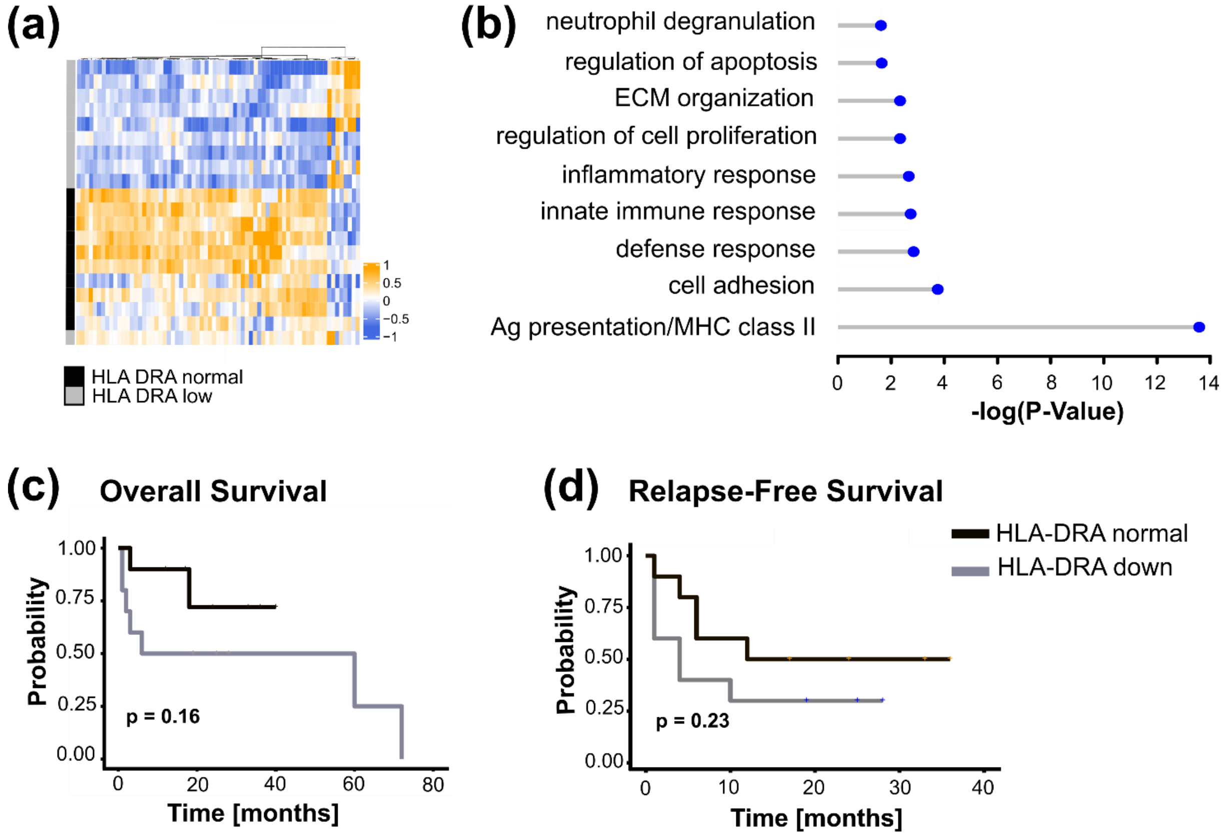


**Supplementary Fig. 2 *HLA-DRA* gene expression was not associated with cytotoxic T cells in PCNSL**

**a.** Differentially expressed genes between PCNSL with normal *HLA-DRA* gene expression (light orange) and *HLA-DRA* gene expression down (purple): 69 significantly deregulated genes were selected (FDR <0.01, $log_2$-FC>4 in absolute value, average read count >20), which segregated the two sample clusters perfectly.

**b.** Main GOBP terms found significantly enriched in genes in Fig. S2a (hypergeometric test, FDR <0.05, at least 3 deregulated genes in the GO term).

**c.** Overall Survival of PCNSL patients with respect to *HLA-DRA* gene expression (Kaplan-Meier curves, log-rank test, n = 20, normal = above the median, low = below the median).

**d.** Relapse Free Survival of PCNSL patients with respect to *HLA-DRA* gene expression (Kaplan-Meier curves, log-rank test, n = 20, normal= above the median, low = below the median).

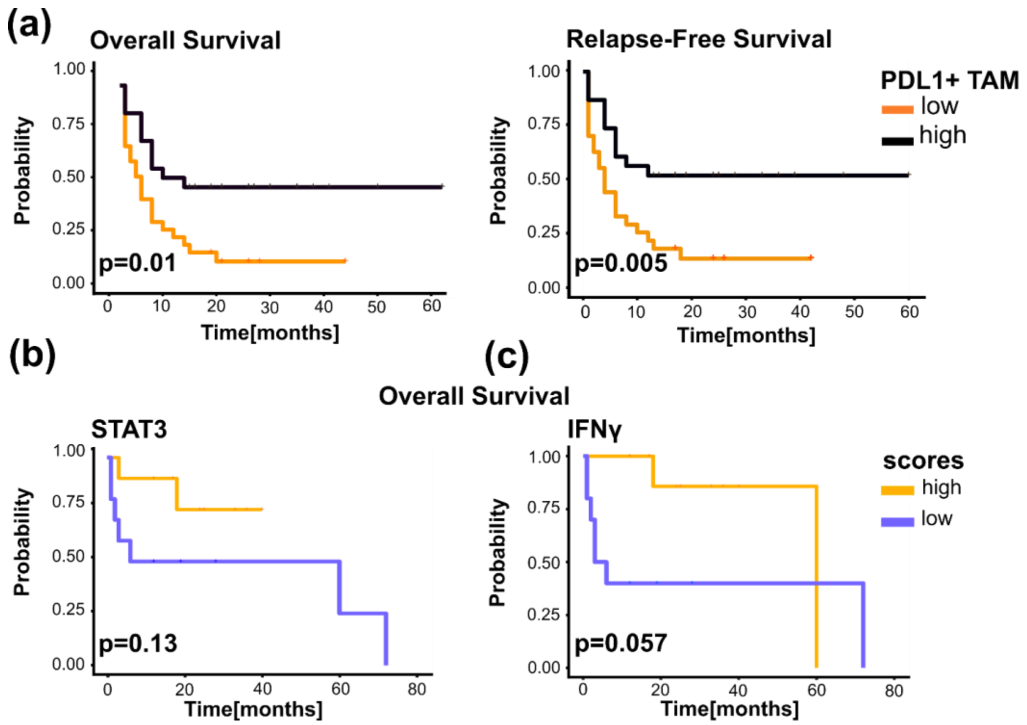


**Supplementary Fig. 3 Immune signaling in PCNSL**

**a.** High PD-L1+TAMs was associated with a better outcome (Kaplan-Meier curves, log-rank test, n = 57). High TAM density (2-3) and a high percentage of CD68+ cells expressing PD-L1 protein (> 50% of CD68+ cells) define high PD-L1+ TAMs.

**b.** Overall Survival of PCNSL patients with respect to STAT3 gene signature score (Kaplan-Meier curves, log-rank test, n = 20, high = above the median, low = below the median).

**c.** Overall Survival of PCNSL patients with respect to IFNγ gene signature score (Kaplan-Meier curves, log-rank test, n = 20, high = above the median, low = below the median).
